# White Matter Microstructural Alterations in Autism Localized with Bundle Analytics

**DOI:** 10.1101/2024.10.16.618731

**Authors:** Gaon S. Kim, Bramsh Q. Chandio, Sebastian M. Benavidez, Paul M. Thompson, Katherine E. Lawrence

## Abstract

Previous diffusion magnetic resonance imaging research in autism has reported altered microstructure in the corpus callosum (CC), the major commissure connecting the two brain hemispheres. However, the CC consists of many fibers connecting various brain regions, and the precise localization of these differences in autism remains unclear. Additionally, the conventional diffusion tensor imaging (DTI) model cannot resolve complex fiber configurations and may miss subtle alterations. In this study, we investigated CC microstructure in 365 participants (5-25 years; 34.0% female; 195 with autism) from 10 sites/scanners by using a novel along-tract mapping method, BUndle ANalytics (BUAN), and advanced microstructure model, the tensor distribution function (TDF). Autism was associated with regionally specific reductions in white matter integrity and elevations in diffusivity. The TDF model detected CC microstructure differences more sensitively than DTI. Taken together, BUAN and TDF allow for precise and rigorous white matter analyses, enhancing the detection of subtle neural differences.

## 1. INTRODUCTION

Autism is a heterogeneous neurodevelopmental condition that has been consistently linked to white matter alterations in the brain using diffusion magnetic resonance imaging (dMRI) [1-2]. Much of the prior dMRI research in autism has used the traditional reconstruction model, diffusion tensor imaging (DTI), together with conventional analytic methods, such as voxel-based analysis (VBA), tract-based spatial statistics (TBSS), or tractography - typically averaging signals across large portions of a given tract [2-4] Many such studies report lower white matter integrity in autism, but the affected white matter tracts are inconsistently identified across studies, as is the localization of such differences within white matter regions [2-4]. These inconsistent results may be due to low statistical power, small effect sizes, modest sample sizes, as well as methodological limitations such as failure to account for crossing fibers, registration errors, and limited anatomical specificity [5-7].

The advanced tractometry method, BUndle ANalytics (BUAN), addresses several limitations of conventional analytic methods by extracting individual white matter tracts and conducting a spatially specific analysis along the entire length of each tract [8-9]. Additionally, the advanced tensor distribution function (TDF) model addresses the limitations of DTI by fitting a continuous mixture of tensors at every voxel, modeling complex fiber orientations [6-7].

Here, we examined white matter microstructure differences in autism in a relatively large sample of 365 participants from 10 sites/scanners using the tractometry method, BUAN, and the TDF microstructure model [6-8] along with DTI microstructural metrics. We focused on the corpus callosum, as prior autism studies have repeatedly implicated this structure but the precise localization and regional specificity of microstructural differences within the corpus callosum remains poorly understood.

## 2. METHODS

### 2.1. Participants and MRI Acquisition

Our final sample consisted of 365 participants between the ages of 5 to 25 years (age: 13.6 ± 3.6 years; 34.0% female). This included 195 autistic individuals (age: 13.4 ± 3.7 years; 34.4% female) and 170 neurotypical controls (age: 13.9 ± 3.3 years; 33.5% female). Data in this sample were drawn from 10 sites/scanners, including seven from the NIMH Data Archive (NDA) and three from the Autism Brain Imaging Data Exchange (ABIDE). More details on participant recruitment, study inclusion criteria, site-specific scanner descriptions, and scanning protocols are detailed elsewhere [10-13]. Briefly, all brain dMRI scans were single-shell acquisitions collected on 3T scanners. The protocol consisted of one or more *b* = 0 s/mm^2^ volume and diffusion-weighted volumes with *b*-values of 1000 s/mm^2^ (8 scanners), 1500 s/mm^2^ (1 scanner), and 2500 s/mm^2^ (1 scanner). The number of diffusion directions varied by site: 32 directions (1 scanner), 48 (1 scanner), 61 (2 scanners), and 64 (6 scanners). To be included in the current analyses, participants from each site were required to have a complete dMRI scan and complete information on diagnosis, age, sex, and full-scale intelligence quotient (FSIQ). Subjects were excluded for poor quality dMRI data (see *Data Analysis*). As we were specifically interested in the developmental period from childhood through emerging adulthood, participants were excluded if they were over 24 years old. For sites that included any siblings or any longitudinal scans, we selected one sibling or time-point per family or participant to maintain the independence of data points. When contrasting the autism and neurotypical groups in our final sample, there were no significant group differences in age (p=0.11) or sex (p=0.86). As expected, FSIQ was higher in neurotypical participants, on average (p<0.001). We thus included FSIQ as a covariate in our analysis (see *Statistical Analysis*).

### 2.2. Data Analysis

The dMRI scans were preprocessed using the ENIGMA DTI protocol [14]. Briefly, preprocessing included eddy current correction, bias field correction, and susceptibility artifact correction, with denoising applied only when appropriate (e.g., yielding randomly distributed residuals, improving signal-to-noise ratio). After preprocessing, we fit the diffusion models, DTI and TDF. DTI, the conventional modeling approach for dMRI data, fits a single tensor to dMRI data and typically represents hindered or isotropic diffusion [5, 15]. However, DTI has well-known limitations in capturing complex fiber configurations [7, 15]. Some of these limitations are addressed by the TDF model developed by our group – a more advanced single-shell model [6-7, 15]. TDF uses a continuous mixture of tensors to account for multiple underlying fiber populations [6]. For each voxel, we calculated four DTI metrics – fractional anisotropy (FA^DTI^), mean diffusivity (MD), axial diffusivity (AD), radial diffusivity (RD) – and one TDF metric, a TDF-derived measure of fractional anisotropy (FA^TDF^) [5-6, 15]. Among DTI metrics, FA^DTI^ measures the degree of anisotropy and MD represents the overall magnitude of water diffusion [1, 16]. AD measures diffusion along the primary axis of the tract and RD measures diffusion perpendicular to it [16]. The TDF metric, FA^TDF^ measures the degree of directional diffusion as does FA^DTI^ – but accounts for crossing fibers.

We analyzed white matter microstructure in autism at a fine-grained anatomical scale using the tractography approach, BUAN (**Fig. 1**) [8]. Briefly, BUAN can precisely localize diagnostic effects along the length of a tract by dividing the tract into many individual segments, based on a parametrized set of streamlines. BUAN captures white matter bundle profiles more accurately than traditional tractometry methods by assigning a segment number to each point on a streamline based on the closest Euclidean distance to a model bundle’s centroid streamline. Unlike some conventional methods, BUAN does not simplify a large white matter bundle to a single mean streamline, but ensures the entire bundle’s microstructural profile is captured [17].

**Fig 1.**
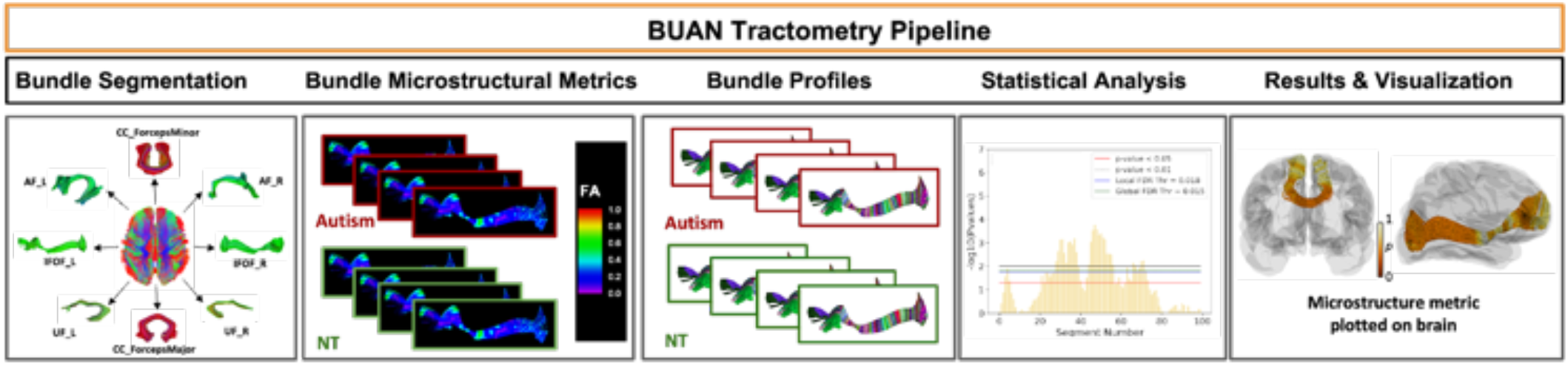
Overview of the BUAN tractometry pipeline. Individual white matter tracts are extracted from a reconstructed whole-brain tractogram. Bundle profiles are created from the extracted bundles by dividing each white matter tract into 100 segments. Next, bundle profiles are analyzed using linear mixed models. *P*-values for the effect of diagnosis are then plotted from these statistical analyses for each microstructural metric at each segment; multiple *p*-value thresholds for significant differences are shown for completeness, with corrections for the false discovery rate (FDR) across the number of tracts and across the number of segments. These *p*-values are mapped onto the tract in a 3D glass brain for visualization.

As in our prior work developing and applying BUAN, whole brain tractograms were generated by using the constrained spherical deconvolution (CSD) model and local deterministic tractography [8, 17-18]. The following parameters were used for fiber tracking: fiber tracking started from voxels with FA values over 0.3, seed count of 10, continuing with a step size of 0.5 mm, an angle curvature threshold of 25°, and tracking stopped at a minimum FA value of 0.2. Streamline-based linear registration (SLR) was used to align individual whole-brain tractograms with the MNI 152 template [8, 19]. Next, RecoBundles was used to extract individual white matter tracts for each participant using their whole brain tractogram and model bundles from the HCP-842 template included in BUAN [8, 20-21]. A tract profile for each bundle was obtained, and the DTI and TDF metrics were mapped onto all the points of the streamlines in a bundle for each subject [8, 22]. Lastly, BUAN was used to divide each tract into 100 segments along the length of the tract.

To ensure data quality, we completed rigorous quality control throughout the processing pipeline [23]. All raw dMRI scans were visually inspected for quality assurance to exclude scans with artifacts. After each step in the dMRI processing pipeline, we visually inspected 2D images of the processing output. Whole-brain tractograms underwent detailed visual inspection for any site/scanner that displayed a low whole-brain streamline count, and the distribution of tract missingness was inspected per site/scanner. Here, we focused on 3 commissural tracts that were reported in the original BUAN report [8]: the *forceps minor, forceps major*, and midbody of the corpus callosum.

### 2.3. Statistical Analysis

Linear mixed models (LMMs) were used to examine differences between the autism and neurotypical groups for each tract segment and metric. Specifically, each white matter metric was set as the dependent variable in the LMM. Diagnosis, age, de-meaned age squared, sex, and FSIQ were included as fixed effects; we focused on the effect of diagnosis, with the other variables included as covariates. As in our prior work, subject and site/scanner were modeled as nested random effects [8, 17]; the former was included to account for the non-independence of different streamlines within a single tract segment for a given subject, and the latter was included to account for potential effects of site/scanner [8, 17, 24].

Correction for multiple comparisons was conducted by controlling the false discovery rate (FDR) [25]. Similar to our prior work, we visualize results at the following multiple *p*-value thresholds for completeness and to allow for comparability with other studies: uncorrected *p* < 0.05, uncorrected *p* < 0.01, with FDR correction across the number of tracts (global FDR threshold), and with FDR correction across the number of segments within each tract (local FDR threshold) [8, 17]. We focus on results obtained at the most stringent FDR threshold, to provide additional rigor [17]. Quantile-Quantile (QQ) plots were created by pooling and visualizing the *p-*values of each metric across the 3 tracts included here [26].

## 3. RESULTS

Microstructural alterations in autism were detected in localized regions of all three commissural tracts examined (**Fig. 2-4**). The *forceps minor, forceps major*, and midbody of the corpus callosum all had lower FA^DTI^ and higher MD and RD in the autism group compared to the neurotypical group. No significant differences in AD were observed between groups. Compared to neurotypical controls, FA^TDF^ was lower for all three tracts in autism. Across metrics, the significant microstructure differences observed in the *forceps minor* and midbody of the corpus callosum were specifically localized to the medial portion of the tract (**Fig. 2-3**). On the other hand, significant differences in the *forceps major* were found in many localized segments across the tract, particularly in the lateral portions (**Fig. 4**). When considering metrics derived from our two distinct microstructure models, DTI and TDF, FA^TDF^ exhibited similar or greater statistical significance than all DTI metrics across all three commissural tracts (**Fig. 5**)

**Fig 2.**
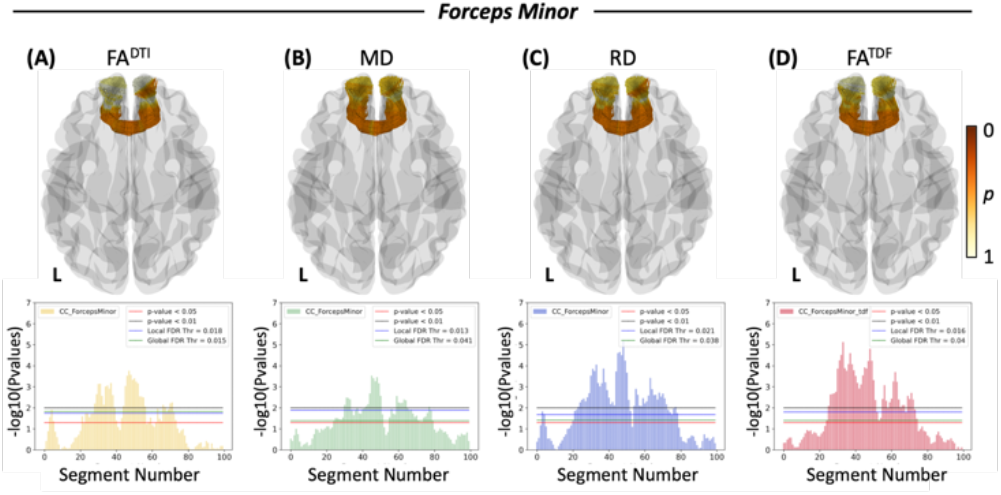
White matter microstructural alterations in *forceps minor* in autism compared to neurotypical controls. Each column represents a different white matter metric: (A) FA^DTI^, (B) MD, (C) RD, and (D) FA^TDF^. The first row is a 3D representation of the *forceps minor* illustrating the anatomical location and corresponding *p*-values of the corresponding microstructural metric. L indicates the left brain hemisphere and the color bar shows the *p*-values, where darker orange is a lower *p*-value nearing 0 and white is a *p*-value of 1. The second row depicts the negative logarithms of *p*-values for each segment along the tract when contrasting the autism and neurotypical groups. FA^DTI^, fractional anisotropy calculated from the diffusion tensor imaging model; MD, mean diffusivity; RD, radial diffusivity; FA^TDF^, fractional anisotropy as calculated by the tensor distribution function model.

**Fig 3.**
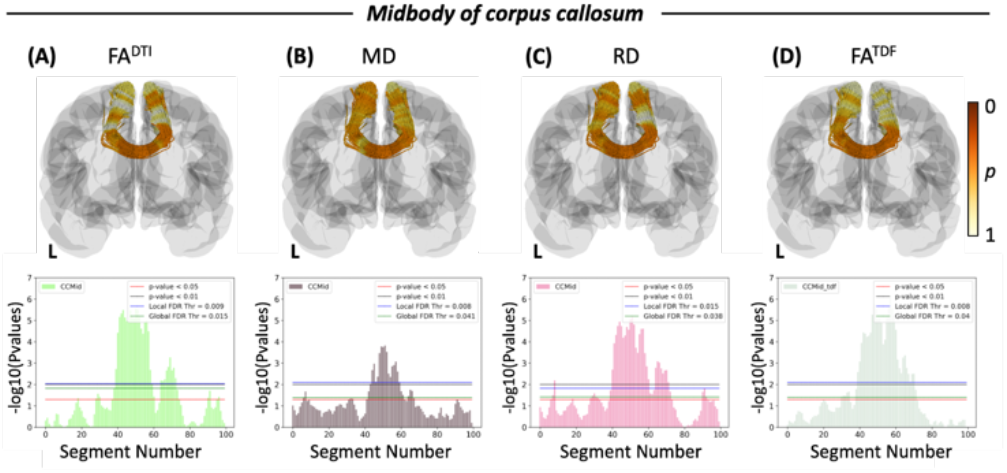
White matter microstructural alterations in the midbody of the corpus callosum in autism compared to neurotypical controls. Each column represents a different white matter metric: (A) FA^DTI^, (B) MD, (C) RD, and (D) FA^TDF^. The first row is a 3D representation of the corpus callosum midbody illustrating the anatomical location and corresponding *p*-values of the corresponding microstructural metric. Abbreviations and color scheme are as in Fig. 2

**Fig 4.**
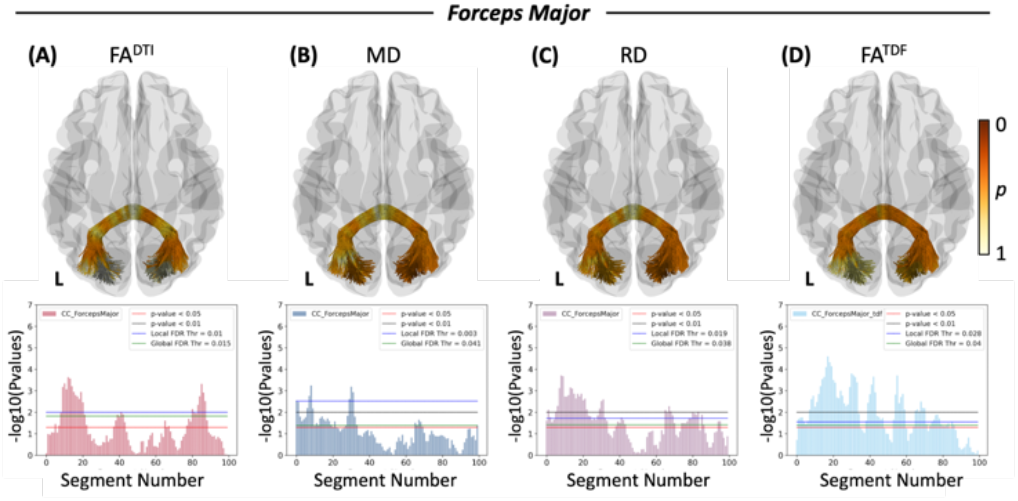
White matter microstructural alterations in *forceps major* in autism compared to neurotypical controls. Each column represents a different white matter metric: (A) FA^DTI^, (B) MD, (C) RD, and (D) FA^TDF^. The first row is a 3D representation of the *forceps major* illustrating the anatomical location and corresponding *p*-values of the corresponding microstructural metric. Abbreviations and color scheme are as in Fig. 2.

**Fig 5.**
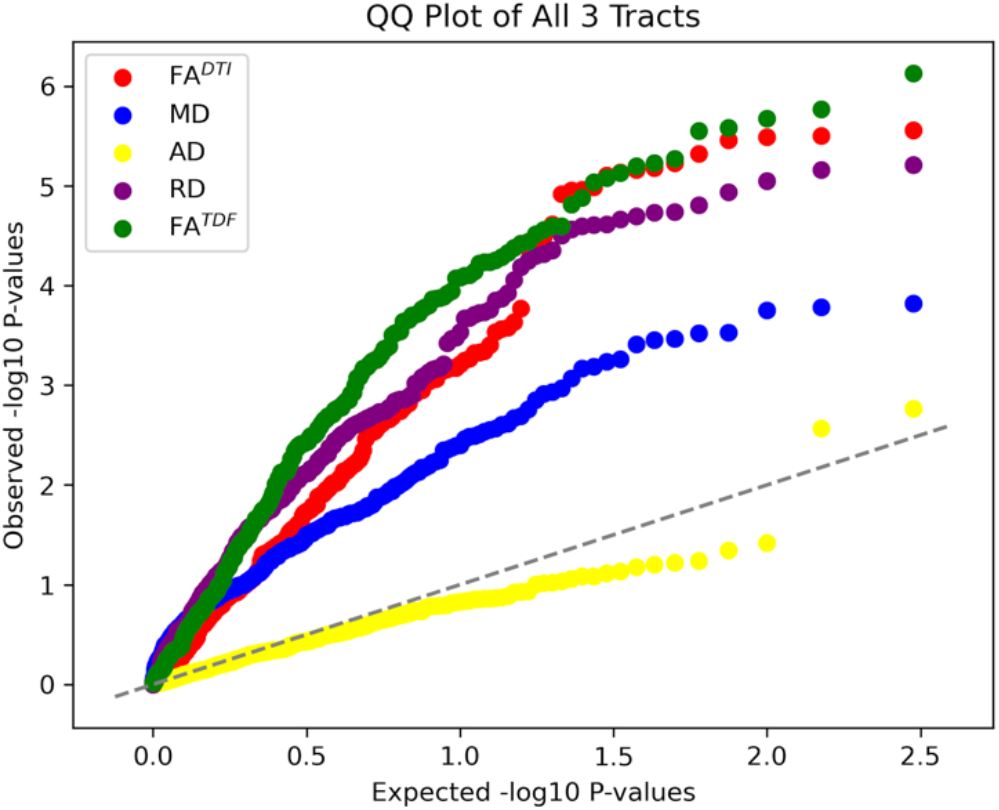
Quantile-Quantile plot of each microstructural metric. This plot shows the *p-*values by each microstructural metric across all 3 corpus callosum tracts. Abbreviations for metrics are as in Fig. 2.

## 4. DISCUSSION

In this study, we used the along-tract analysis framework, BUAN, and the advanced single-shell TDF to localize white matter microstructural alterations in a large-scale sample of autistic and neurotypical individuals. All three commissural tracts examined here exhibited lower white matter FA (often a sign of reduced integrity) and higher diffusivity in autism in spatially specific segments along the tract, demonstrating that commissural white matter alterations in autism are localized within individual tracts but not confined to a specific area or lobe of the brain.

Our study has several strengths. Our use of BUAN provided finer-scale spatial localization of microstructural differences in autism, expanding on prior work implicating the corpus callosum in autism [1-4]. We also used the TDF microstructural model, which addresses the well-known limitations of DTI by capturing complex fiber orientations. TDF overall improved sensitivity to microstructural alterations in autism, enhancing sensitivity to subtle neural differences. Future studies should extend these findings by examining additional white matter tracts, including association, projection, and cerebellar tracts [8]. Such future work should also examine multi-shell diffusion MRI data to examine additional microstructural metrics, such as those provided by the neurite orientation dispersion and density imaging (NODDI) model or diffusion spectrum imaging (DSI), to provide further insight into white matter microstructural differences in autism [27].

In sum, our findings in a relatively large sample of 365 participants with and without autism indicate that autism is associated with significant white matter alterations across commissural tracts. These alterations are localized within specific segments of these tracts rather than uniformly affecting the entire tract. The dMRI methods BUAN and TDF captured these microstructural differences in a spatially precise and sensitive manner, demonstrating the promise of these methods for advancing research in autism and other brain-based conditions.

## 5. COMPLIANCE WITH ETHICAL STANDARDS

This research study was conducted retrospectively using human subject data made available in open access by (NIMH Data Repositories and ABIDE II team). These analyses were approved by the Institutional Review Board of the University of Southern California.

## 6. ACKNOWLEDGMENTS

This research was supported by the National Institutes of Health (award RF1AG057892 to P.M.T.), the Momental Foundation (Mistletoe Research Fellowship to K.E.L.), and the Asan Foundation (Biomedical Science Scholarship to G.S.K.). Research reported in this publication was also supported by the Office Of The Director, National Institutes Of Health of the National Institutes of Health under Award Number S10OD032285.

Data used in the preparation of this article reside in part in the NIH-supported NIMH Data Repositories, a collaborative informatics system created by the NIH to provide a national resource to support and accelerate research in mental health related conditions. Dataset identifiers: nda2021 (6 sites/scanners) and nda1906 (1 site/scanner). We thank the ABIDE II team for providing data (at http://fcon_1000.projects.nitrc.org/indi/abide). Dataset identifiers: Trinity College Dublin (1 site/scanner), San Diego State University (1 site/scanner), Barrow Neurological Institute (1 site/scanner). This manuscript reflects the views of the authors and does not reflect the opinions or views of the NIH or of the Submitters submitting original data to the NIMH Data Repositories or ABIDE II.

## REFERENCES

[1] B. G. Travers et al., “Diffusion Tensor Imaging in Autism Spectrum Disorder: A Review,” Autism Research, vol. 5, no. 5, pp. 289–313, Oct. 2012, doi: 10.1002/aur.1243.

[2] Y. Zhao, L. Yang, G. Gong, Q. Cao, and J. Liu, “Identify aberrant white matter microstructure in ASD, ADHD and other neurodevelopmental disorders: A meta-analysis of diffusion tensor imaging studies,” Progress in Neuro-Psychopharmacology and Biological Psychiatry, vol. 113, p. 110477, Mar. 2022, doi: 10.1016/j.pnpbp.2021.110477.

[3] X. Di, A. Azeez, X. Li, E. Haque, and B. B. Biswal, “Disrupted focal white matter integrity in autism spectrum disorder: A voxel-based meta-analysis of diffusion tensor imaging studies,” Progress (i) in Neuro-Psychopharmacology and Biological Psychiatry, vol. 82, pp. 242–248, Mar. 2018, doi: 10.1016/j.pnpbp.2017.11.007.

[4] Y. Aoki, O. Abe, Y. Nippashi, and H. Yamasue, “Comparison of white matter integrity between autism spectrum disorder subjects and typically developing individuals: a meta-analysis of diffusion tensor imaging tractography studies,” Mol Autism, vol. 4, no. 1, p. 25, 2013, doi: 10.1186/2040-2392-4-25.

[5] P. J. Basser, J. Mattiello, and D. LeBihan, “MR diffusion tensor spectroscopy and imaging,” Biophysical Journal, vol. 66, no. 1, pp. 259–267, Jan. 1994, doi: 10.1016/S0006-3495(94)80775-1.

[6] T. M. Nir et al., “Fractional anisotropy derived from the diffusion tensor distribution function boosts power to detect Alzheimer’s disease deficits,” Magnetic Resonance in Medicine, vol. 78, no. 6, pp. 2322–2333, 2017, doi: 10.1002/mrm.26623.

[7] A. D. Leow et al., “The tensor distribution function,” Magnetic Resonance in Med, vol. 61, no. 1, pp. 205–214, Jan. 2009, doi: 10.1002/mrm.21852.

[8] B. Q. Chandio et al., “Bundle analytics, a computational framework for investigating the shapes and profiles of brain pathways across populations,” Sci Rep, vol. 10, no. 1, p. 17149, Oct. 2020, doi: 10.1038/s41598-020-74054-4.

[9] E. Garyfallidis et al., “Dipy, a library for the analysis of diffusion MRI data,” Front. Neuroinform., vol. 8, Feb. 2014, doi: 10.3389/fninf.2014.00008.

[10] J. E. Lee et al., “Diffusion tensor imaging of white matter in the superior temporal gyrus and temporal stem in autism,” Neurosci Lett, vol. 424, no. 2, pp. 127–132, Sep. 2007, doi: 10.1016/j.neulet.2007.07.042.

[11] A. Di Martino et al., “Enhancing studies of the connectome in autism using the autism brain imaging data exchange II,” Sci Data, vol. 4, p. 170010, Mar. 2017, doi: 10.1038/sdata.2017.10.

[12] A. Irimia, C. M. Torgerson, Z. J. Jacokes, and J. D. Van Horn, “The connectomes of males and females with autism spectrum disorder have significantly different white matter connectivity densities,” Sci Rep, vol. 7, no. 1, p. 46401, Apr. 2017, doi: 10.1038/srep46401.

[13] A. Jack et al., “A neurogenetic analysis of female autism,” Brain, vol. 144, no. 6, pp. 1911–1926, Jul. 2021, doi: 10.1093/brain/awab064.

[14] N. Jahanshad et al., “Multi-site genetic analysis of diffusion images and voxelwise heritability analysis: A pilot project of the ENIGMA–DTI working group,” NeuroImage, vol. 81, pp. 455–469, Nov. 2013, doi: 10.1016/j.neuroimage.2013.04.061.

[15] D. Jones, “Studying connections in the living human brain with diffusion MRI,” Cortex, vol. 44, no. 8, pp. 936–952, Sep. 2008, doi: 10.1016/j.cortex.2008.05.002.

[16] J. M. Soares, P. Marques, V. Alves, and N. Sousa, “A hitchhiker’s guide to diffusion tensor imaging,” Front. Neurosci., vol. 7, 2013, doi: 10.3389/fnins.2013.00031.

[17] L. Nabulsi et al., “Multi-Site Statistical Mapping of Along-Tract Microstructural Abnormalities in Bipolar Disorder with Diffusion MRI Tractometry,” bioRxiv, p. 2023.08.17.553762, Oct. 2023, doi: 10.1101/2023.08.17.553762.

[18] B. Jeurissen, J.-D. Tournier, T. Dhollander, A. Connelly, and J. Sijbers, “Multi-tissue constrained spherical deconvolution for improved analysis of multi-shell diffusion MRI data,” NeuroImage, vol. 103, pp. 411–426, Dec. 2014, doi: 10.1016/j.neuroimage.2014.07.061.

[19] E. Garyfallidis, O. Ocegueda, D. Wassermann, and M. Descoteaux, “Robust and efficient linear registration of white-matter fascicles in the space of streamlines,” NeuroImage, vol. 117, pp. 124–140, Aug. 2015, doi: 10.1016/j.neuroimage.2015.05.016.

[20] E. Garyfallidis et al., “Recognition of white matter bundles using local and global streamline-based registration and clustering,” Neuroimage, vol. 170, pp. 283–295, Apr. 2018, doi: 10.1016/j.neuroimage.2017.07.015.

[21] F.-C. Yeh et al., “Population-averaged atlas of the macroscale human structural connectome and its network topology,” NeuroImage, vol. 178, pp. 57–68, Sep. 2018, doi: 10.1016/j.neuroimage.2018.05.027.

[22] E. Garyfallidis, M. Brett, M. M. Correia, G. B. Williams, and I. Nimmo-Smith, “QuickBundles, a Method for Tractography Simplification,” Front Neurosci, vol. 6, p. 175, Dec. 2012, doi: 10.3389/fnins.2012.00175.

[23] S. M. Benavidez et al., “Sex Differences in the Brain’s White Matter Microstructure during Development assessed using Advanced Diffusion MRI Models,” bioRxiv, p. 2024.02.02.578712, 2024, doi: 10.1101/2024.02.02.578712.

[24] B. Q. Chandio et al., “Bundle Analytics based Data Harmonization for Multi-Site Diffusion MRI Tractometry,” May 01, 2024, bioRxiv. doi: 10.1101/2024.02.03.578764.

[25] Y. Benjamini and Y. Hochberg, “Controlling the False Discovery Rate: A Practical and Powerful Approach to Multiple Testing,” Journal of the Royal Statistical Society: Series B (Methodological), vol. 57, no. 1, pp. 289–300, Jan. 1995, doi: 10.1111/j.2517-6161.1995.tb02031.x.

[26] B. Q. Chandio et al., “Amyloid, Tau, and APOE in Alzheimer’s Disease: Impact on White Matter Tracts,” bioRxiv, p. 2024.08.05.606560, Aug. 2024, doi: 10.1101/2024.08.05.606560.

[27] H. Zhang, T. Schneider, C. A. Wheeler-Kingshott, and D. C. Alexander, “NODDI: Practical in vivo neurite orientation dispersion and density imaging of the human brain,” NeuroImage, vol. 61, no. 4, pp. 1000–1016, Jul. 2012, doi: 10.1016/j.neuroimage.2012.03.072.

